# BIOTIA-DX RESISTANCE Achieved the Best Antimicrobial Resistance Phenotype Prediction Accuracy at CAMDA 2025

**DOI:** 10.1101/2025.09.24.678344

**Authors:** Gábor Fidler, Heather Wells, Ford Combs, John Papciak, Mara Couto-Rodriguez, Sol Rey, Tiara Rivera, Lorenzo Uccellini, Christopher E. Mason, Niamh B. O’Hara, Dorottya Nagy-Szakal, David C. Danko

**Affiliations:** Biotia Inc., New York, NY, USA; Tri-Institutional Computational Biology & Medicine Program, Weill Cornell Medicine of Cornell University, New York, NY, USA; The HRH Prince Alwaleed Bin Talal Bin Abdulaziz Alsaud Institute for Computational Biomedicine, Weill Cornell Medicine, New York, NY, USA; The WorldQuant Initiative for Quantitative Prediction, Weill Cornell Medicine, New York, NY, USA; The Feil Family Brain and Mind Research Institute, Weill Cornell Medicine, New York, NY, USA; SUNY Downstate Health Sciences University, The Department of Cell Biology, College of Medicine, New York, NY, USA

## Abstract

We have developed BIOTIA-DX RESISTANCE (BDXR), a bioinformatic tool for predicting antimicrobial resistance (AMR) from whole genome sequencing of microbial isolates. BDXR achieved the best accuracy of any submission to the CAMDA 2025 AMR Challenge. This year’s challenge focused on predicting AMR **phenotype** which is a more complex problem than the detection of AMR marker genes, the focus of some prior years. BDXR achieved an overall F1 score of 89% on the training set and 84.1% on the challenge test set. Accuracy varied across the 9 species and drug pairs in the competition from an F1 score of 98.4% (*Campylobacter jejuni*, tetracycline) to 78.5% (*Pseudomonas aeruginosa*, cefatzidime). BDXR is based on curation of global datasets, machine learning-based predictions from input data, and highly stringent prepreprocessing of input data and databases.

## Background

### Antimicrobial Resistance

Antimicrobial resistance (AMR) is a global threat projected to become a leading cause of death by 2050 (Kariuki 2024). Persistent selective pressure from antimicrobial overuse has driven the adaptation of resistant strains. Genetic causes of AMR are highly diverse. Resistance can be due to the presence of a single gene, variations within a gene, or interactions between multiple gene variants. Common resistance pathways include drug inactivation due to enzymatic activity, drug target alteration and active efflux.

While drug inactivation is often caused by acquisition of a single gene, in many cases complex gene network interactions are the underlying source of resistance (Reygaert 2018). Moreover, distribution of antimicrobial resistance genes (ARGs) vary in different bacteria and their expression may be influenced by several factors. The gene families involved may be clade- and drug-specific or may have broad range and effect. This diversity means that there are a huge number of genetic factors involved in drug resistance with exponentially more potentially meaningful combinations.

Efforts to catalogue ARGs through standard microbiological workflows have produced meaningful databases of genes, particularly for clinically relevant organisms. These databases represent gold standards with clear causal attribution between specific genetic mechanisms and resistance but leave substantial gaps where microbes display resistance in phenotypic testing without displaying known genetic markers of resistance. The number of potential factors involved in drug resistance necessitates computational approaches that can make effective predictions from more diffuse databases.

While there are relatively few data that provide causal links between specific genes and AMR, there are substantially more datasets that link microbial genomes with susceptibility data. These data lack clear genetic attribution. While the genomes in the database certainly contain genes that result in drug resistance, it is not clear which genes, variants, or combinations specifically result in the resistance. Further, these data are not fully curated. While organized databases like PATRIC exist, a large number of microbial genomic data are stored only in public nucleotide archives and the text of publications. Effective use of broad global data sources is crucial to develop more predictive models of AMR from genetic data.

### Phenotype Prediction vs Gene Markers

There are a variety of tools that detect the presence of known ARGs. Though related, this is a different simpler problem than predicting AMR phenotype from genomic data. Most relevant species-drug pairs have multiple mechanisms by which a microbe can become resistant to that drug. While some gene markers do strongly correlate with a specific phenotype it is often the case that phenotypes may have multiple potential causes. As an example, fluoroquinolone resistance in *E. coli* can be attributed to specific mutations in either gyrA or in parC (in addition to other potential causes, Jacoby 2005). In these cases the absence of a marker is insufficient to rule out the phenotype. Additionally, there are many cases where resistance phenotype is determined by multiple genetic markers.

### The CAMDA AMR Competition

The Critical Assessment of Massive Data Analysis (CAMDA) is an annual set of competitions for prediction techniques relevant to human health (Łabaj et al. 2023). One of the annual constituent competitions is CAMDA AMR, a competition to predict antimicrobial resistance from microbial genomic data. The CAMDA AMR 2025 competition consisted of predicting resistant phenotype for 5,345 genomes split across 9 species drug pairs. This included predictions for four drugs (gentamicin, erythromycin, tetracycline, and cefatzidime) and included both gram positive and negative species. The competition consists of a training set of public data with known resistance phenotype and a test set of newly sequenced data where the resistance phenotype is sequestered.

## Results

### Performance on the CAMDA AMR Training and Test Sets

BIOTIA-DX Resistance achieved an F1 score of 84.1% on the CAMDA test set and an overall F1 score of 89% on the CAMDA training set. The breakdown of F1 score by species and drug on the training set can be found in table 1. We did not have access to a category breakdown for the test set.

**Table 1:**
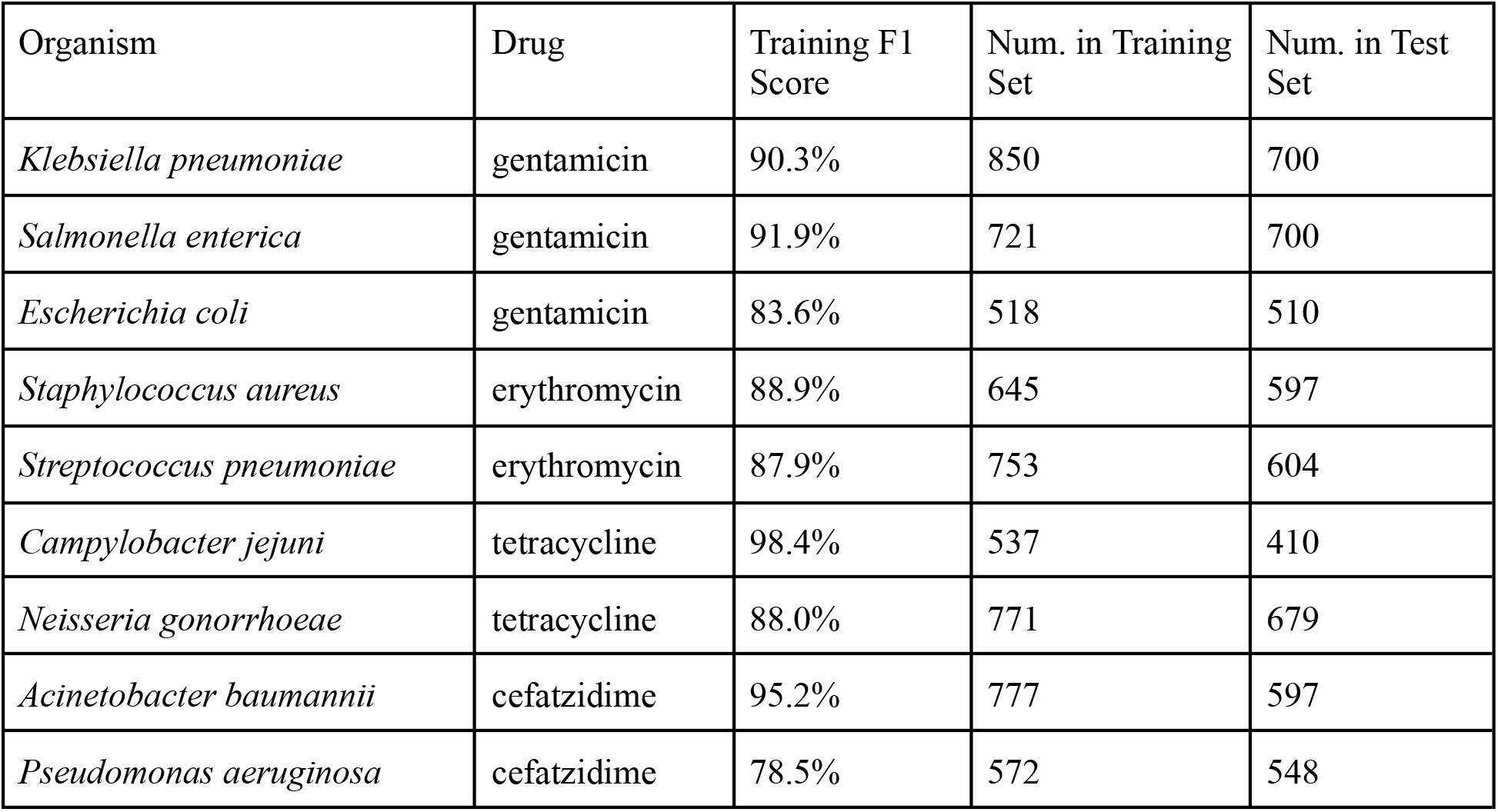
The training set accuracy of BDXR on different species drug pairs. F1 scores are the mean of 5-fold cross validation.

On the training set the F1 score ranged from 98% for *Campylobacter jejuni* (tetracycline) down to 78% for *Pseudomonas aeruginosa* (ceftazidime), with an average score of 89%. The F1 score was above 85% for all but two species drug pairs. *P. aeruginosa* (ceftazidime) and *E. coli* (gentamicin) were less accurate, possibly due to multigenic resistance patterns or a larger number of relevant genetic variants. F1 scores on the training set were obtained via cross-validation and as such do not represent memoization of the training set.

### Performance relative to other submissions to CAMDA AMR

Table 2 lists the top five leaderboard scores by group at the time of competition close (May 15, 2025). Where groups had multiple submissions we show the best score achieved. Three groups including BIOTIA-DX Resistance achieved F1 scores over 80% on the test set.

**Table 2:** Best scores achieved by the top 5 groups by the competition deadline.

| Group | Leaderboard Test Score |
| --- | --- |
| BIOTIA-DX Resistance | 84 |
| Davuluri Lab | 82 |
| Yurtseven & Djamalova | 82 |
| CCM - fsnieto | 78 |
| weijiang | 55 |

## Methods

### Datasets

CAMDA AMR provided two related datasets: a training set and a test set. The training set consisted of microbial genome sequencing data and paired resistance phenotype information. Resistance phenotype was given as an ordered categorical variable with *susceptible, intermediate*, and *resistant* values. We treated *intermediate* values as *resistant*. Minimum inhibitory concentration (MIC) values were also given for a subset of samples, but we did not make use of this information. The microbial genome sequencing data was variable. All samples were based on whole genome sequencing but depth varied as did sequencing type (e.g. short read, long read).

The test set consisted of microbial sequencing data only. The resistance phenotypes were known to the CAMDA organizers for each genome but this information was sequestered. As with the training set the sequencing data consisted of whole genome sequencing but with variable sequencing methodology and depth. The organizers of CAMDA AMR affirmed that phenotypes for the test data did not appear in any public repository. We did not have access to the phenotype data for the test set.

### Training and Evaluation

To train BDXR we collected over 10,000 genomic sequences with known phenotypes from the supplied CAMDA training data, the Bacterial and Viral Bioinformatics Resource Center (BV-BRC), the Comprehensive Antibiotic Resistance Database (CARD) and other publicly available sources (Olson et al. 2023), (Alcock et al. 2023). Data was curated on our platform GeoSeeq. We identified known AMR genes in these genomes and generated clusters of genetically similar genes. Gene clusters were identified based on Jaccard similarity of k-mer spectra.

Each genome was described as a vector of gene presence or absence. For gene families where we determined variants to be relevant, we selected variants to include in the vector by filtering to only those which were statistically significantly associated with the resistant phenotype. Vectors were subject to unsupervised dimensionality reduction. The resulting vectors were paired with susceptibility data as labels and used to develop predictive models.

Using the input vectors described above, we trained a family of machine learning models for taxonomic and drug resistance groups. Model input was filtered based on feature importance to reduce dimensionality and training was subject to cross-validation to reduce overfitting. Once features were selected, models were subject to hyperparameter tuning and an optimal prediction probability threshold was selected based on F1 score. Model selection was performed by training multiple models and selecting the best performer. Training F1 accuracy scores were obtained by averaging 5-fold cross validation data.

After training we generated phenotype predictions for the CAMDA test set. These predictions were submitted to the CAMDA leaderboard which automatically generated an overall score. Competition rules allowed up to three submissions to the test set and we submitted twice.

## Discussion

Prediction of AMR from microbial genomic data remains an important unsolved medical problem. There are numerous clinical situations where traditional antimicrobial susceptibility testing (AST) is infeasible due to factors like uncultivability, expected turn around time, low biomass of samples, sample type and more. In many of these cases sequencing is already a beneficial technology for its ability to identify a very wide range of microbial species. The ability to detect drug resistance from sequencing data greatly enhances the ability of clinicians to provide effective targeted care.

Clinically validated diagnostics to identify the presence of relevant ARG marker genes already exist. However, this is a simpler problem than predicting AMR phenotype. While clinically useful marker gene identification cannot rule out resistance caused by an alternate mechanism nor can it detect polygenic resistance. True AMR phenotype prediction would greatly enhance the clinical utility of molecular microbial testing by allowing clinicians to make more informed decisions.

As CAMDA demonstrates, effective prediction of complex resistance phenotype from sequencing data is possible. However, prediction accuracy of many species drug pairs remains below desirable clinical accuracies. The central difficulty of resistance prediction is the wide and evolving variety of mechanisms for resistance. Large datasets are critical to capturing a wide variety of potential mechanisms. Ideally these datasets would consist of both in vitro AST as well as real world efficacy results showing how patients responded to different prescribed drugs.

Further improvements in resistance prediction accuracy will also require improved models that can effectively use large datasets. Deep learning architectures are still only lightly used in biology but growing datasets, practical success in other fields, and new microbiology foundation models are likely to drive a substantial increase in the number and effectiveness of deep learning techniques (Zhou et al. 2025; Brixi et al. 2025). Our group is developing deep learning approaches to AMR phenotype prediction as are other groups that submitted to CAMDA.

Though deep learning approaches are likely to lead to practical improvements in prediction accuracy, the research community should continue to investigate novel AMR mechanisms. Many deep learning tools lack effective explanatory abilities and research into AMR mechanisms is likely to remain critical to the development of future antibiotic drugs.

## Data Availability

The CAMDA AMR Leaderboard is publicly available on the CAMDA website at https://bipress.boku.ac.at/camda2025/competitions/camda-2025/?leaderboard.

